# Citrullination of pyruvate kinase by PADI1 and PADI3 regulates glycolysis and cancer cell proliferation

**DOI:** 10.1101/718486

**Authors:** Sebastien Coassolo, Guillaume Davidson, Luc Negroni, Giovanni Gambi, Sylvain Daujat, Christophe Romier, Irwin Davidson

**Author notes:** To whom correspondence should be addressed E mail.

## Abstract

CHD3 and CHD4 are mutually exclusive ATPase subunits of the Nucleosome Remodelling and Deacetylation (NuRD) complex that regulates gene expression. CHD4 is essential for growth of multiple patient derived melanoma xenografts and for breast cancer. Here we show that CHD4 regulates expression of PADI1 (Protein Arginine Deiminase 1) and PADI3 in multiple cancer cell types modulating citrullination of three arginines of the allosterically-regulated glycolytic enzyme pyruvate kinase M2 (PKM2). Citrullination reprograms cross-talk between PKM2 ligands lowering its sensitivity to the inhibitors Tryptophan, Alanine and Phenylalanine and promoting activation by Serine. Citrullination thus bypasses normal physiological regulation by low Serine levels to promote excessive glycolysis defining a novel pathway regulating proliferation of melanoma and other cancer cells. We provide unique insight as to how conversion of arginines to citrulline impacts key interactions within PKM2 that act in concert to reprogram its activity as an additional mechanism regulating this important enzyme.

## Introduction

A hallmark of cancer cells is the high glycolysis and lactic acid production under aerobic conditions, a metabolic state known as the Warburg effect ^1^. Tumour tissues accumulate increased amounts of glucose used not only for energy production, but also for anabolic reactions. Glycolytic intermediates are notably used for *de novo* synthesis of nucleotides or amino acids like glycine and serine produced in large amounts to sustain high rates of cancer cell proliferation ^2, 3^. Coupling of energy production via glycolysis to the availability of the intermediates required for nucleotide and amino acid synthesis is controlled in large part by an alternatively spliced isoform of the enzyme pyruvate kinase called PKM2 expressed in proliferating embryonic and cancer cells ^4, 5^. Unlike the PKM1 isoform that is constitutively active, PKM2 activity is positively regulated by serine (Ser), fructose 1,6-biphosphate (FBP) an intermediate of the glycolytic pathway and succinylaminoimidazole-carboxamide riboside (SAICAR), an intermediate in *de novo* purine nucleotide synthesis ^4, 6, 7^. High levels of these molecules stimulate PKM2, but when their levels are lowered through excessive glycolysis, PKM2 activity is inhibited by amino acids such as tryptophan (Trp), alanine (Ala) and phenylalanine (Phe) that compete with Ser to allosterically regulate PKM2 activity ^8–10^. Through this complex feedback loop, PKM2 couples glycolytic flux to the level of critical intermediate metabolites. PKM2 activity is also regulated by post-translational modifications, such as tyrosine phosphorylation or proline hydroxylation under hypoxia ^11, 12^.

Melanoma cells are no exception to the Warburg effect, showing high levels of aerobic glycolysis induced by transformation with oncogenic BRAF or NRAS ^13^. Treatment with vemurafenib, an inhibitor of oncogenic BRAF, down-regulates aerobic glycolysis, regained in resistant cells ^14^. Transcription factor MITF (Microphthalmia associated transcription factor) regulates many parameters of melanoma cell physiology including metabolism ^15^. MITF directly regulates PPARGC1 and cells with high MITF expression show elevated oxidative phosphorylation compared to cells with low MITF with higher glycolysis ^16, 17^.

Bossi et al ^18^ performed an shRNA dropout screen to identify proteins involved in epigenetics and chromatin functions essential for patient derived melanoma xenograft (PDX) growth. This screen identified the ATPase subunit of the PBAF chromatin remodelling complex BRG1 along with CHD3 and CHD4, the catalytic ATPase subunits of the Nuclesome Remodelling and Deacetylation (NuRD) complex, as essential for tumour formation by all tested melanoma PDX. NuRD, is an epigenetic regulator of gene expression, acting in many, but not all ^19^, contexts as a co-repressor that remodels chromatin through its ATPase subunits and deacetylates nucleosomes through its HDAC1 and HDAC2 subunits ^20–23^. CHD4 has also been reported to be essential in breast cancer ^24^. Here, we describe a novel pathway where CHD4 regulates expression of the PADI1 (Protein Arginine Deiminase 1) and PADI3 enzymes that convert arginine to citrulline. Increased PADI1 and PADI3 expression enhances citrullination of three arginines of the key glycolytic enzyme PKM2 leading to excessive glycolysis, lowered ATP levels and slowed cell growth. CHD4 therefore links epigenetic regulation of PADI1 and PAD3 expression to glycolytic flux and the control of cancer cell growth.

## Results

### CHD4 regulates PADI1 and PADI3 expression in melanoma cells

We performed siRNA CHD3 and CHD4 silencing in a collection of melanoma cells. Silencing was specific for each subunit as measured by RT-qPCR and confirmed by immunoblot (Fig. 1a-b). Loss of CHD3 also mildly reduced MITF expression, whereas that of SOX10 was unchanged. In agreement with the results of the previous shRNA dropout screen, siRNA-mediated CHD3 or CHD4 silencing reduced clonogenic capacity, increased the proportion of slow proliferating cells (Fig. 1a-d), but did not induce apoptosis (Fig. 1e). The effects were less dramatic than seen upon silencing of MITF that induces a potent cell cycle arrest and senescence ^25, 26^. RNA-seq following CHD4 silencing in melanoma cells identified more than 1000 up-regulated genes compared to 364 down-regulated genes showing that CHD4 was primarily acting as a transcriptional repressor (Fig. 1f-g, and Supplementary Dataset 1). In contrast, similar numbers of genes were up or down-regulated by CHD3 silencing (Fig. 1f-g and Supplementary Dataset 1), but no significant overlap between the CHD3 and CHD4 up and down-regulated genes and the pathways they regulate were observed (Fig. 1h-i). These data accord with previous reports showing that CHD3 and CHD4 form NuRD complexes with mainly distinct functions ^27^. De-regulated expression of selected genes upon CHD4 silencing was confirmed by RT-qPCR on independent RNA samples in both 501Mel and MM117 melanoma cells (Fig. 1J-K).

**Figure 1.**
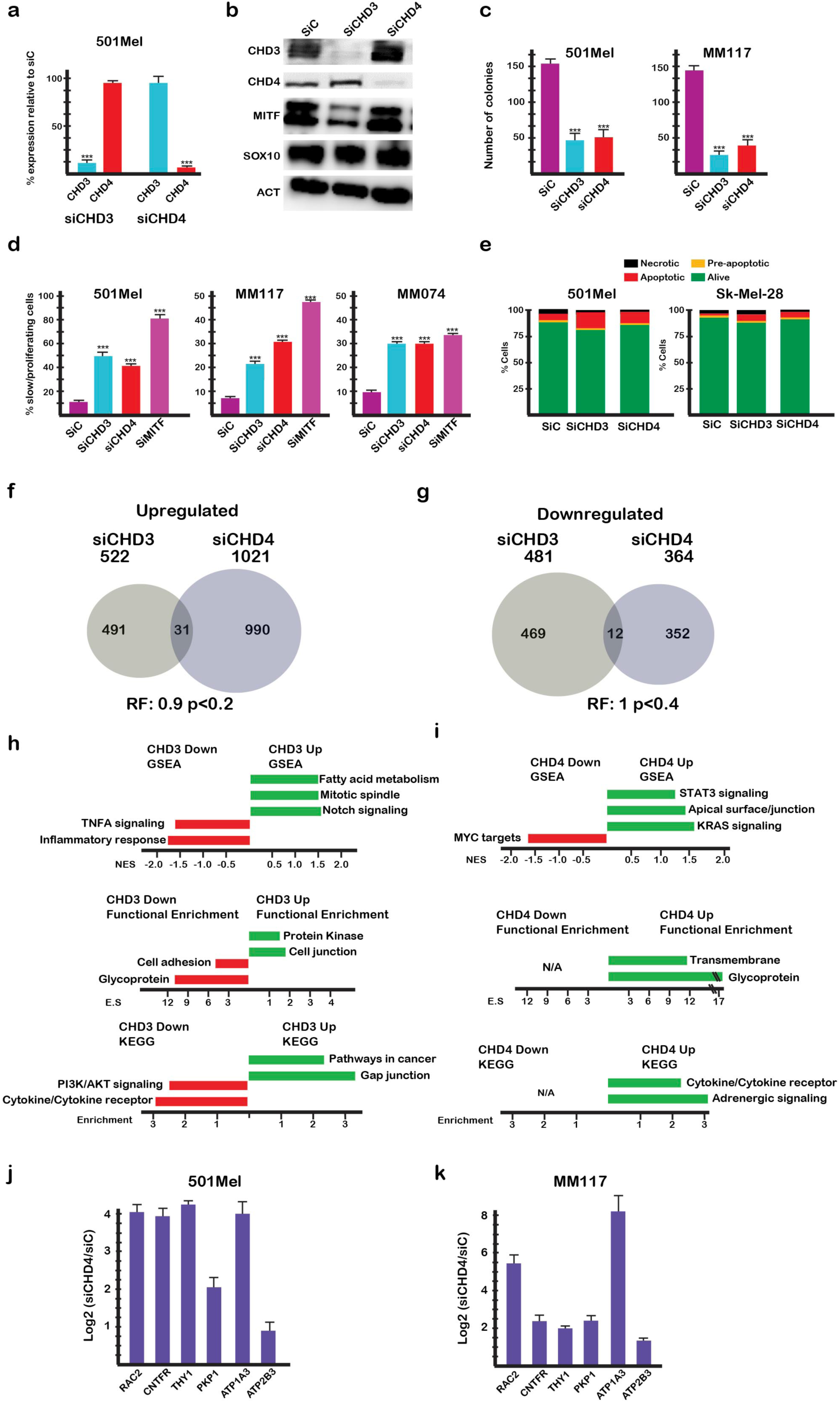
CHD3 and CHD4 are required for normal melanoma cell proliferation. **a-b**. 501Mel cells were transfected with the indicated siRNAs and CHD3 and CHD4 expression evaluated by RT-qPCR or by immunoblot along with that of MITF and SOX10. **c**. The indicated cell lines were transfected with siRNA and after reseeding the number of colonies counted after 10 days. **d**. The indicated cell lines were transfected with siRNAs and cell proliferation evaluated by cell trace violet assay. **e**. The indicated cell lines were transfected with siRNA and apoptosis detected by FACs after labelling with Annexin-V. Silencing of MITF known to induce cell cycle arrest and senescence was included as a control. **f-g**. CHD3 and CHD4 regulate distinct gene expression programs and functional pathways. Genes up or down-regulated based on Log2 fold-change >1/<-1 with an adjusted p-value <0,05 were identified. Venn diagrams show overlap between the CHD3 and CHD4 regulated genes along with the hypergeometric probability representation factor (RF), in this case non-significant. **h-i.** Ontology analyses of CHD3 and CHD4 regulated genes. Shown are the enrichment scores for GSEA, as well as David functional enrichment and KEGG pathway categories. **j-k**. Verification of deregulated expression of selected genes following siCHD4 in independent RNA samples from 501Mel or MM117 cells. In all experiments N=3 and unpaired t-test analyses were performed by Prism 5. P-values: *= p<0,05; **= p<0,01; *\*\*\*= p<0,001.* Data are mean ± SEM.

Amongst the genes potently up-regulated by CHD4, but not CHD3, silencing are *PADI1* (Protein Arginine Deiminase 1) and *PADI3* encoding enzymes that convert arginine to citrulline ^28^ (Fig. 1j-k, Fig. 2a-b). In all tested melanoma lines, *PADI3* expression was almost undetectable and potently activated by CHD4 silencing, whereas some others had low basal *PADI1* levels also strongly stimulated by CHD4 silencing (Fig. 1h). RNA-seq further showed that expression of *PADI2* and *PADI4* was low to undetectable and their expression was not induced by siCHD4 silencing (Supplementary Dataset 1).

**Figure 2.**
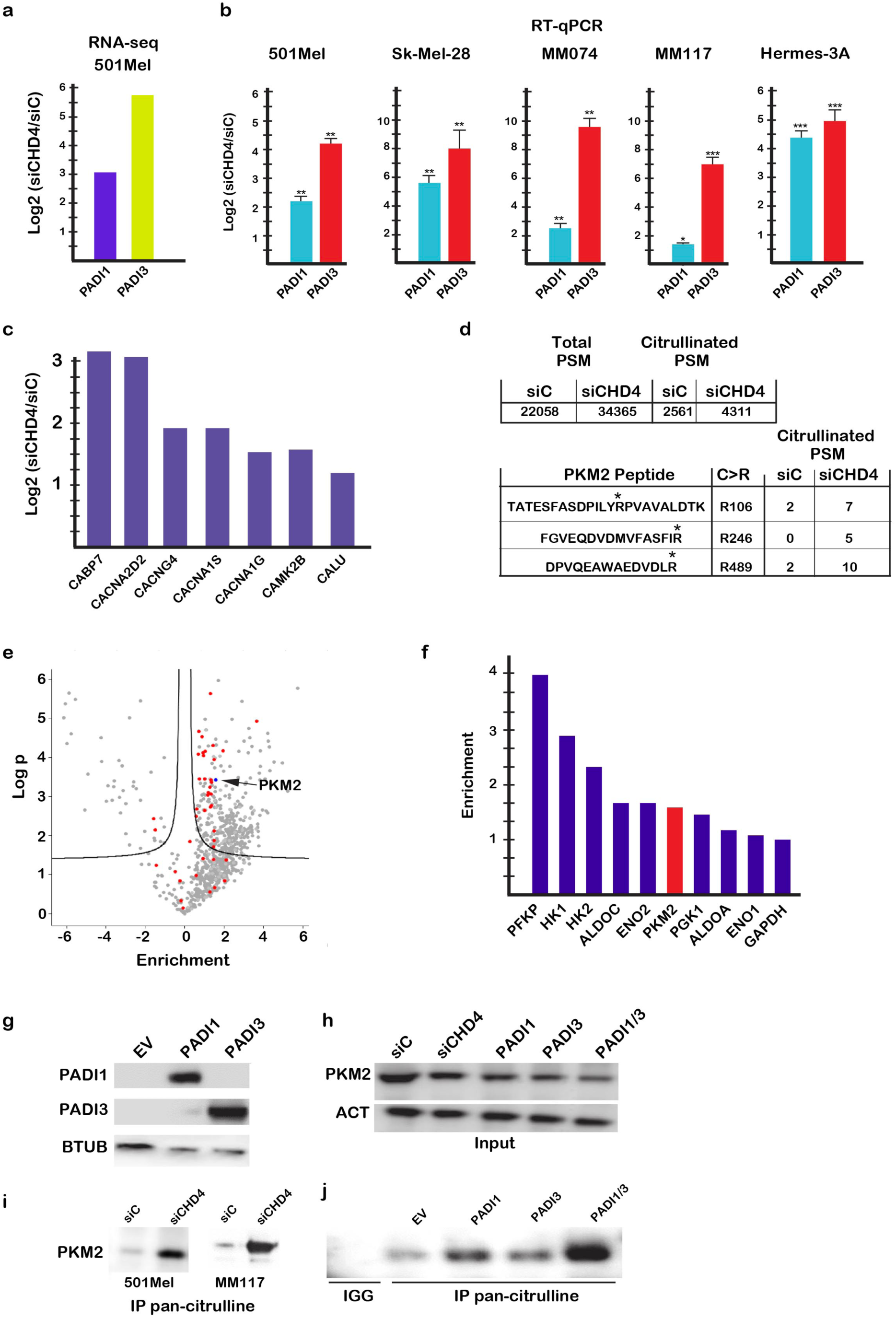
CHD4 regulates *PADI1* and *PADI3* expression and citrullination of their substrates. **a-b**. *PADI1* and *PADI3* expression in the indicated cells lines following CHD4 silencing shown by RNA-seq (A) and RT-qPCR (B). **c.** Increased expression of gene involved in calcium signaling following CHD4 silencing. **d.** CHD4 silencing increases citrullination in 501Mel cells. Increases in number of total and citrullinated PSMs in CHD4 silenced cells following immunoprecipitation (IP) with pan-citrulline antibody. Lower table shows PKM2 peptides with increased citrullination after pan-citrulline IP. **e**. Volcano plot showing proteins with increased or decreased total PSMs after pan-citrulline IP. **f**. Increased recovery of glycolytic enzymes following pan-citrulline IP. **g.** Immunoblot showing expression of recombinant PADI1 and PADI3 in cells transfected with the corresponding expression vectors of the empty vector (EV). **h.** Immunoblot showing expression of PKM2 in cells after CHD4 silencing or transfection with the PADI1 and PADI3 vectors in the cell extracts used for immunoprecipitation with pan-citrulline antibody. **i.** PKM2 in the pan-citrulline IPs from 501Mel or MM117 cells. **j.** Immunoblot showing PKM2 in the pan-citrulline IP after transfection with the PADI1 and/or PADI3 expression vectors.

Arginine citrullination by PADI enzymes is a calcium-dependent reaction. CHD4 silencing also strongly up-regulated a set of genes involved in calcium signalling including calcium channel subunits, calcium binding proteins and calcium-dependent protein kinases (Fig. 2c) consistent with CHD4 silencing inducing PADI enzyme expression and activity.

The *PADI1* and *PADI3* genes are located next to each other (Supplementary Fig. 1). ChIP-seq in melanoma cells revealed that CHD4 occupied an intronic regulatory element in *PADI1* immediately adjacent to sites occupied by transcription factors CTCF and FOSL2 (AP1). This element is predicted to regulate both the *PADI1* and *PADI3* genes (Supplementary Fig. 1) and is further marked by H2AZ, H3K4me1, BRG1 and ATAC-seq, but not by the lineage-specific transcription factors MITF and SOX10. CHD4 therefore appears to repress the activity of this element to prevent activation by CTCF, AP1 or other transcription factor that may bind to it.

Analyses of the Cancer Cell Line Encyclopaedia showed a strong correlation of *PADI1* and *PADI3* expression indicating their co-regulation was not restricted to melanoma cells (Supplementary Fig. 2a). Remarkably, expression of both *PADI1* and *PADI3* was negatively correlated with both *CTCF* and *CHD4* (Supplementary Fig. 2b-e), but was positively correlated with that of *FOSL1/2* and *JUNB*. These data support the idea that CHD4-CTCF act to repress the locus that is activated by AP1 (Supplementary Fig. 2f-i).

Analyses of TCGA human tumour datasets showed strongly positively correlated expression of *PADI1* and *PADI3* in multiple tumour types such as melanoma, uveal melanoma, bladder, lung and head and neck, with *PADI1* often being the gene showing strongest correlation with *PADI3* (Supplementary Fig. 3a-f). Despite this strong co-regulation, no negative correlation with *CHD4* or *CTCF* was seen with the exception of uveal melanoma, where *PADI1/3* expression was negatively correlated with *CTCF*, but positively correlated with *FOSL1* and *JUNB* analogous to what was seen in cancer cell lines (Supplementary Fig. 3g-i). As *CTCF* and *CHD4* are highly and rather ubiquitously expressed, insufficient enrichment in tumour cells in the TCGA samples may have masked their negative correlation with *PADI1* and *PADI3*. Alternatively, *PADI1* and *PADI3* co-expression tumours may be regulated by other mechanisms. Nevertheless, these data indicated that *PADI1* and *PADI3* were co-ordinately regulated in multiple cancer cell types and human tumours.

### PADI1 and PADI3 citrullinate glycolytic enzymes and stimulate glycolysis

To identify potential PADI1 and PADI3 substrates in melanoma cells, we made protein extracts from siC and siCHD4 cells, performed immunoprecipitation (IP) with a pan-citrulline antibody and analysed precipitated proteins by mass-spectrometry (Fig. 2d and Supplementary Dataset 2). An increased number of total peptide spectral matches (PSMs) and PSMs for citrullinated peptides were detected following CHD4 silencing. A set of predominantly cytoplasmic proteins including tubulins, multiple 14-3-3 proteins and glycolytic enzymes PFKP, HK1/2, GAPDH, ALDOA/C, ENO1/2 and PKM2 were enriched in the pan-citrulline IP from siCHD4 cells (Fig. 2d-f and Supplementary Dataset 2).

We focussed on PKM2, a highly regulated enzyme playing a central role in integrating control of glycolysis with cellular metabolic status and cell cycle ^29^. PKM2 converts phosphoenolpyruvate (PEP) to pyruvate then converted to lactic acid. To investigate PKM2 citrullination, melanoma cells were transfected with siC, siCHD4 or vectors allowing ectopic expression of PADI1 and PADI3 (Fig. 2g). While CHD4 silencing or PADI1/3 expression did not alter overall PKM2 levels (Fig. 2h), strongly increased amounts of PKM2 were detected in the pan-citrulline IP following siCHD4 compared to siC in both 501Mel and MM117 melanoma cells and after ectopic PADI1 and PADI3 expression, particularly upon co-expression of both enzymes (Fig. 2i-j).

We generated antibodies against synthetic peptides corresponding to citrullinated R106 and R246. The R489 peptide was too hydrophobic to obtain soluble peptide. In dot-blot assays, each of these antibodies showed strong signal for the citrullinated peptide, but little for the equivalent wild-type peptide with arginine (Supplementary Fig. 4a-b). Similarly, citrullinated R106 antibody did not recognise citrullinated R246 and vice-versa (Supplementary Fig. 4c-d). Immunoblots on extracts from cells transfected with siCHD4 or PADI1 and PADI3 expression vectors showed enhanced signal for PKM2 compared to the control transfected cells indicating increased citrullination of these two arginines (Supplementary Fig. 4e-f). Note also that PKM2 was essentially the only protein recognised by these antibodies further indicating their specificity.

To determine if siCHD4 silencing and enhanced PKM2 citrullination altered glycolysis, we profiled melanoma cell metabolism in real time. CHD4 silencing in all tested melanoma lines increased the basal OCR (oxygen consumption rate) and ECAR (extracellular acidification rate), markedly increased maximum OCR and ECAR and decreased the OCR/ECAR ratio due to the increased ECAR values (Fig. 3a-d). ECAR was blocked using 2-deoxy-D-glucose confirming that it was due to increased glycolysis (Fig. 3c). Increased glycolysis and lactic acid production diverts pyruvate from oxidative metabolism a more efficient ATP source. Consequently, excessive glycolysis following CHD4 silencing led to decreased intracellular ATP levels (Fig. 3e).

**Figure 3.**
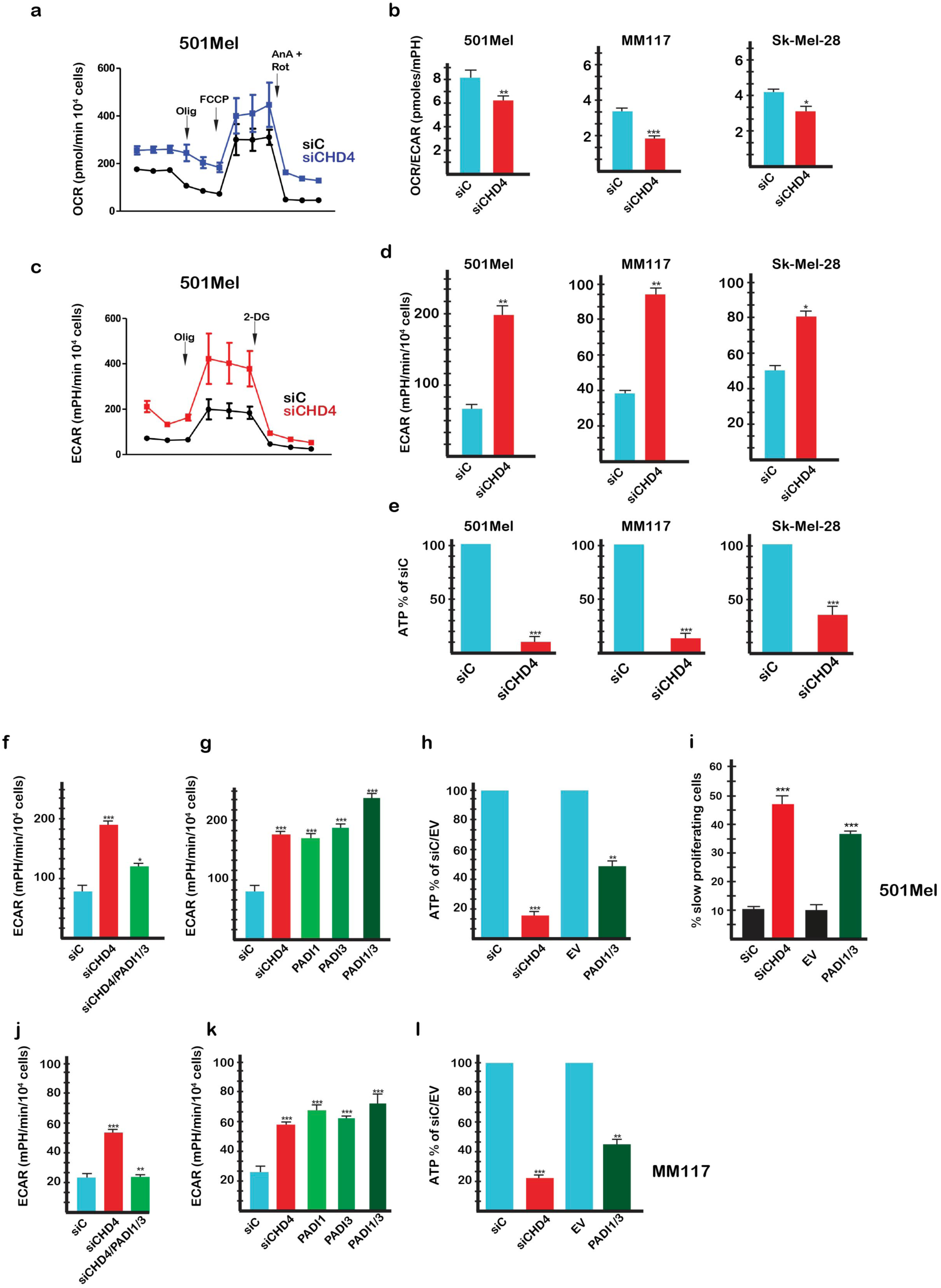
CHD4 silencing regulates glycolysis and cell proliferation. **a.** Effect of CHD4 silencing on basal and maximal OCR values in 501Mel cells. **b**. Effect of CHD4 silencing on the basal OCR/ECAR ratio in the indicated cell types. **c-d**. Effect of CHD4 silencing on basal and maximal ECAR values in 501Mel cells and basal ECAR values in the indicated cell types. **e.** CHD4 silencing reduces intracellular ATP levels in the indicated cell lines. **f-g**. ECAR values in 501Mel cells following transfection with indicated siRNAs or expression vectors. **h.** Intracellular ATP levels following CHD4 silencing or PADI1/3 expression. EV = empty expression vector control. **i.** Reduced cell proliferation following PAD1/3 expression. **j-k**. ECAR values and intracellular ATP levels in MM117 cells following transfection with indicated siRNAs or expression vectors. In all experiments ECAR values were determined from N=6 with 6 technical replicates for each N. Unpaired t-test analysis were performed by Prism. P-values: *= p<0,05; **= p<0,01; ***= p<0,001. Data are mean ± SEM.

The increased glycolysis seen upon CHD4 silencing was strongly diminished when *PADI1* and *PADI3* were additionally silenced (Fig. 3f-j). In contrast, exogenous expression of PADI1, PADI3 or both stimulated glycolysis (Fig. 3g and k). Consistent with increased glycolysis, PADI1/3 expression led to reduced intracellular ATP levels (Fig. 3h-i) and reduced cell proliferation (Fig. 3j). PADI1 and PADI3 were therefore necessary and sufficient for increased glycolysis accounting for the effect seen upon CHD4 silencing.

It has previously been shown that treatment of melanoma cells with BRAF inhibitors induces metabolic reprogramming, strongly reducing glycolysis ^14^. Moreover, dependence on glycolysis sensitizes melanoma cells to the effects of BRAF inhibition ^30^. Consistent with these observations, CHD4 silencing or ectopic PADI1/3 expression that increased glycolysis sensitized Sk-Mel-28 cells to the effects of the BRAF inhibitor vemurafenib (Supplementary Fig. 5). Hence, by regulating glycolysis CHD4 silencing or PADI1 and PADI3 expression acts to modulate melanoma cell sensitivity to BRAF inhibition. Nevertheless, the effect of CHD4 silencing had more potent effects on vemurafenib sensitivity that ectopic PADI1 and PADI3 expression suggesting additional pathways are affected.

### PADI1 and PADI3 stimulate glycolysis in a variety of cancer cell types

As mentioned above, PADI1 and PADI3 were co-ordinately regulated in multiple types of cancers. Moreover, PADI1 and PADI3 expression were positively correlated with that of PKM2 in cancer cells (Supplementary Fig. 2j-k) suggesting that the regulatory mechanism described above in melanoma may be relevant in other cancer types. SiCHD4 silencing in SiHa cervical carcinoma cells strongly diminished their clonogenic capacity (Fig. 4a), potently increased *PADI3* expression (Fig. 4b) and stimulated glycolysis (Fig. 4c-d). Moreover, glycolysis was stimulated by ectopic PADI1/3 expression leading to reduced OCR/ECAR ratio and ATP levels (Fig. 4e-g). In HeLa cells, CHD4 silencing reduced clonogenic capacity and activated *PADI1* and *PADI3* expression (Fig. 4h-i). Glycolysis was stimulated by both CHD4 silencing and ectopic PADI1/3 expression (Fig. 4j). CHD4 silencing strongly stimulated PADI1 and PADI3 expression in MCF7 breast cancer cells and increased glycolysis (Fig. 4k-l). Analogous results were observed in two different types of renal cell carcinoma cell lines, UOK-109 translocation renal cell carcinoma cells (Fig. 4m-q) and A498 clear cell renal carcinoma cells (Fig. 4r-v). Therefore, in cell lines from distinct cancer types, CHD4 silencing or ectopic PADI1/3 expression increased glycolysis and negatively impacted cell proliferation.

**Figure 4.**
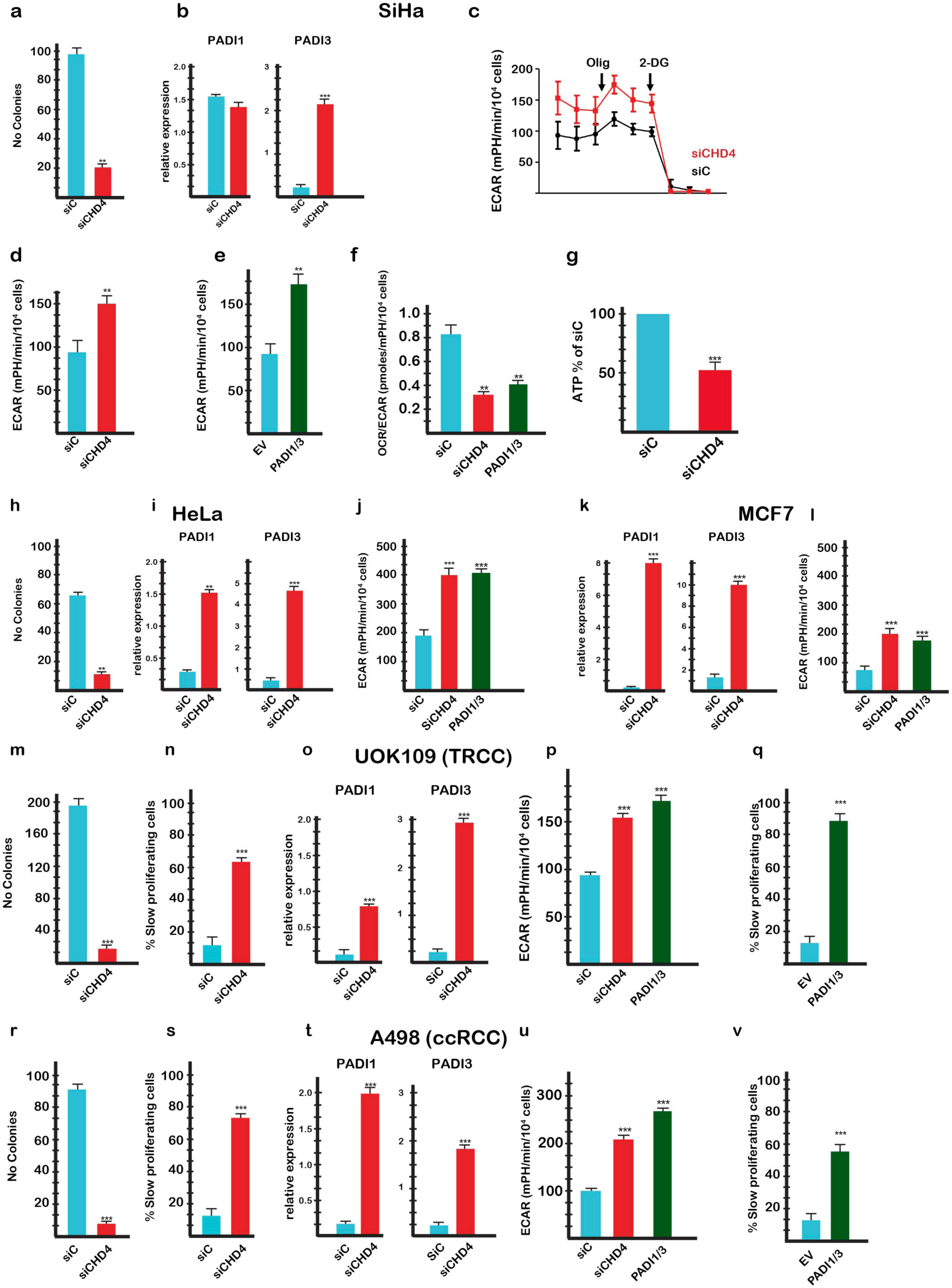
Citrullination regulates glycolysis and proliferation in multiple types of cancer cells. **a**. Diminished clonogenicity of SiHa cells following CHD4 silencing. **b**. *PADI1* and *PADI3* expression in SiHa cells following CHD4 silencing. **c-d**. Basal and maximal glycolysis in SiHa cells following CHD4 silencing. **e.** Glycolysis in SiHa cells following PADI1/3 expression. **f.** OCR/ECAR ratio in SiHa cells following CHD4 silencing or PADI1/3 expression. **g.** Intracellular ATP levels in SiHa cells following CHD4 silencing. **h-j**. Clonogenicity, *PADI1*, *PADI3* expression and glycolysis in HeLa cells following CHD4 silencing or PADI1/3 expression as indicated. **k-l**. *PADI1*, *PADI3* expression and glycolysis in MCF7 cells following CHD4 silencing or PADI1/3 expression as indicated. **m-q.** Clonogenicity, *PADI1, PADI3* expression and glycolysis and proliferation in UOK-109 translocation renal cell carcinoma cells following CHD4 silencing or PADI1/3 expression as indicated. **r-v**. Clonogenicity, *PADI1, PADI3* expression and glycolysis and proliferation of A498 clear cell renal carcinoma cells following CHD4 silencing or PADI1/3 expression as indicated. For RT-qPCR, ATP levels and clonogenicity, N=3 and statistical unpaired t-tests analyses were performed by Prism 5 P-values as above. For glycolysis: N=3 with 6 technical replicates for each N. P-values as above. Data are mean ± SEM.

### Citrullination reprograms PKM2 allosteric regulation

As described in the introduction, PKM2 isoform activity is positively regulated by serine (Ser), and FBP and negatively by Trp, Ala and Phe, thus coupling glycolytic flux to the level of critical intermediate metabolites ^4–6^. PKM2 allosteric regulation involves three distinct enzyme conformations [^8, 9, 31^ and Supplementary Fig. 6a]. In the apo (resting) state, in absence of small molecules and ions, the PKM2 N-terminal and A domains adopt an active conformation, but the B domain is in an inactive conformation. In the activated R-state, binding of FBP or Ser and magnesium, stabilizes the N and A domains in their active conformation, and rotates the B domain towards the A domain that together form the active site. In the inactive T-state, upon binding of inhibitory amino acids (Trp, Ala and Phe), the B domain adopts a partially active conformation, but the N and A domains undergo structural changes and disorganize the active site. The structural changes observed between the different PKM2 states are reinforced allosterically by organisation into a tetramer that is essential for enzyme function.

In siCHD4 extracts, 3 citrullinated arginine residues, R106, R246 and R489 enriched in the siCHD4 extracts were identified by mass-spectrometry and R106 and R246 confirmed by immunoblot (Fig. 2c and Supplementary Fig. 4e-f). In the apo state, R246 forms salt bridges between its guanidino group and the main chain carboxyl groups of V215 and L217 at the pivotal point where the B domain moves between its active and inactive conformations [^31^ and Supplementary Fig. 6b]. This interaction contributes to maintaining the inactive B domain conformation in the apo state and is lost in the R- and T-states. R246 citrullination should strongly weaken or abolish interaction with V215 and L217 facilitating release of the B domain from its inactive conformation.

R106 participates in the free amino acid binding pocket. In the apo state, R106 mostly faces the solvent, but upon free amino acid binding, it rotates towards the pocket where its guanidino group interacts with the carboxylate group of the bound amino acid and the P471 main chain carbonyl [^6, 8, 9^ and Fig. 5a]. Ser forms a hydrogen bond network with the N and A domains stabilizing their active conformations, whereas the hydrophobic side chains of Trp, Ala, or Phe cause displacement of the N-domain outwards leading to the allosteric changes that characterize the inactive T-state (Fig. 5a).

**Figure 5.**
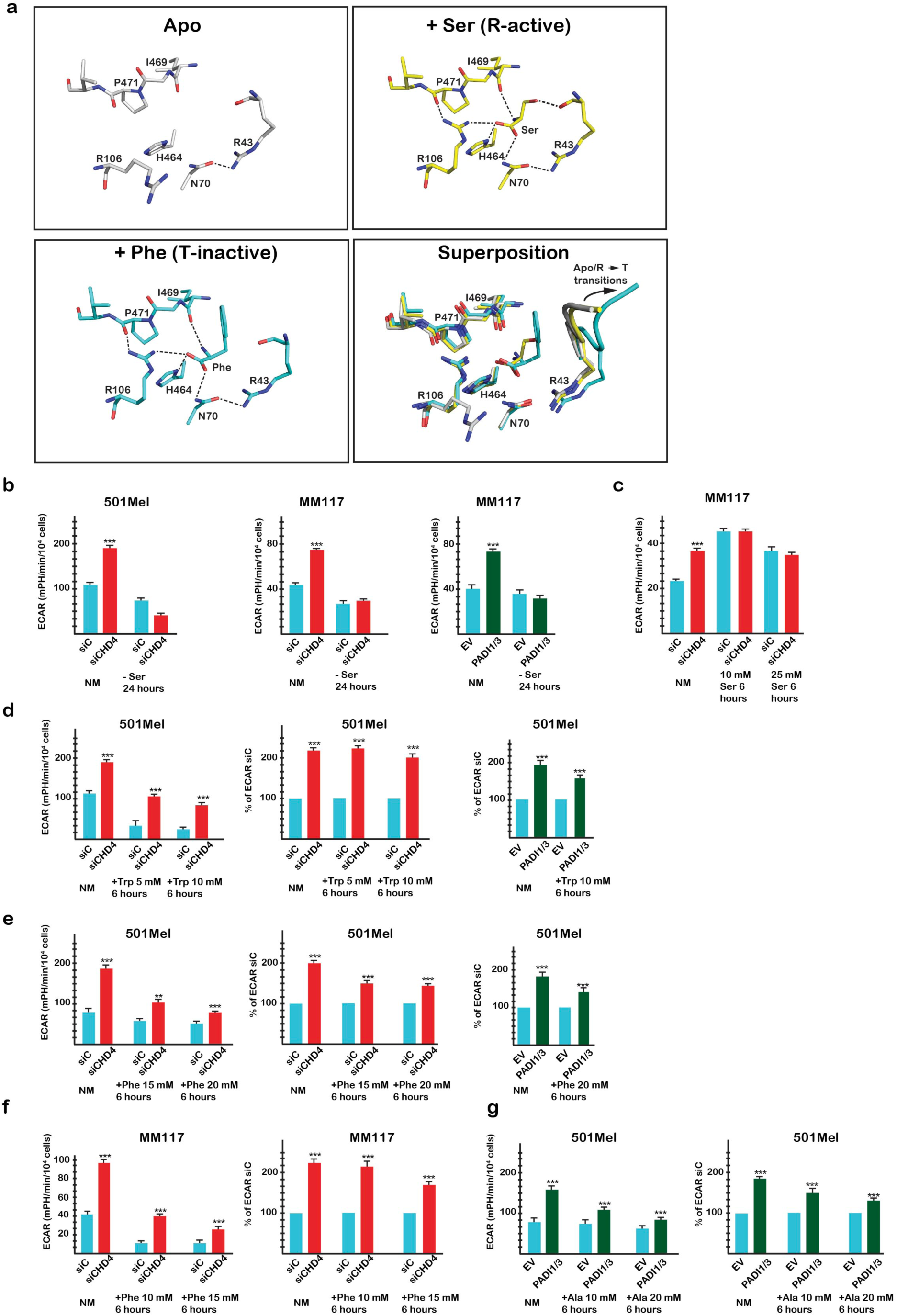
PKM2 citrullination diminishes allosteric inhibition by Phe/Ala/Trp. **a.** Close up view of free Ser and Phe interactions within the free amino acid binding pocket in the Apo, R-active and T-inactive states with a superposition of the three structures. All residues displayed are shown as sticks. In the superposition, the peptide bearing R43 is represented as ribbon to show the allosteric changes created upon Phe binding. Salt bridges and hydrogen bonds are shown as dashed lines. For clarity, the side chain of Phe 470, which stacks on R106 side chain, is not displayed. PDB data sets are as described in Supplementary Figure 3. **b**. ECAR values in absence of Ser after CHD4 silencing or PADI1/3 expression in 501Mel or MM117 cells; NM = normal medium. **c.** ECAR values in presence of exogenous Ser with or without CHD4 silencing in 501Mel cells. **d-e**. ECAR values in presence of exogenous Trp or Phe with or without CHD4 silencing or PADI1/3 expression in 501Mel cells. **f**. ECAR values in presence of exogenous Phe with or without CHD4 silencing in MM117 cells. **g.** ECAR values in presence of exogenous Ala with or without PADI1/3 expression in 501Mel cells. In all experiments ECAR values were determined from N=6 with 6 technical replicates for each N. Unpaired t-test analysis were performed by Prism 5. P-values: *= p<0,05; **= p<0,01; ***= p<0,001. Data are mean ± SEM.

Transition between the R- and T-states is finely regulated by changes in the relative concentrations of Ser versus Trp, Ala and Phe that compete for binding to the pocket ^9^. Loss of R106 positive side chain charge upon citrullination will diminish its ability to interact with the carboxylate group of the free amino acids. Due to its extended network of hydrogen bonds within the pocket and as it does not modify the active conformations of the N and A domains, it is possible that binding of Ser is less affected than that of the hydrophobic amino acids that induce important structural changes within the N and A domains. Consequently, R106 citrullination could reduce the inhibitory effect of Trp, Ala and Phe thereby shifting the equilibrium towards activation by Ser.

To test the above hypotheses, we asked if citrullination modulated glycolysis under different conditions. When cells were grown in absence of Ser, basal glycolysis was reduced and was no longer stimulated upon siCHD4 or PADI1/3 expression (Fig. 5b). On the other hand, exogenous Ser stimulated basal glycolysis to a level that was not further increased by siCHD4 (Fig. 5c). In contrast, basal glycolysis was reduced by exogenous Trp, but remained stimulated by siCHD4 and by PADI1/3 expression (Fig. 5d). Similarly, glycolysis was stimulated by siCHD4 in presence of increasing Phe concentrations (Fig. 5e), an effect particularly visible in MM117 cells where despite strongly inhibited basal glycolysis, stimulation was seen upon siCHD4 (Fig. 5f). PADI1/3 expression also stimulated glycolysis in presence of exogenous Ala (Fig. 5g). PKM2 citrullination did not therefore bypass the requirement for Ser, while excess Ser mimicked stimulation seen by siCHD4. In contrast, siCHD4 or PADI1/3 expression diminished inhibition by Trp/Ala/Phe, consistent with the idea that R106 citrullination modified the equilibrium in favour of the activator Ser.

R489 is directly involved in FBP binding with strong interactions between its guanidino group and the FBP 1’ phosphate group (Fig. 6a). Despite its extensive interaction network with PKM2, FBP binding is severely reduced upon mutation of R489 into alanine ^8, 10^. Hydrogen bonding with R489 therefore plays a critical role in FBP binding that should be diminished by loss of its side chain charge upon citrullination, hence suggesting that activation of PKM2 by citrullination required weakening of its interaction with FBP.

**Figure 6.**
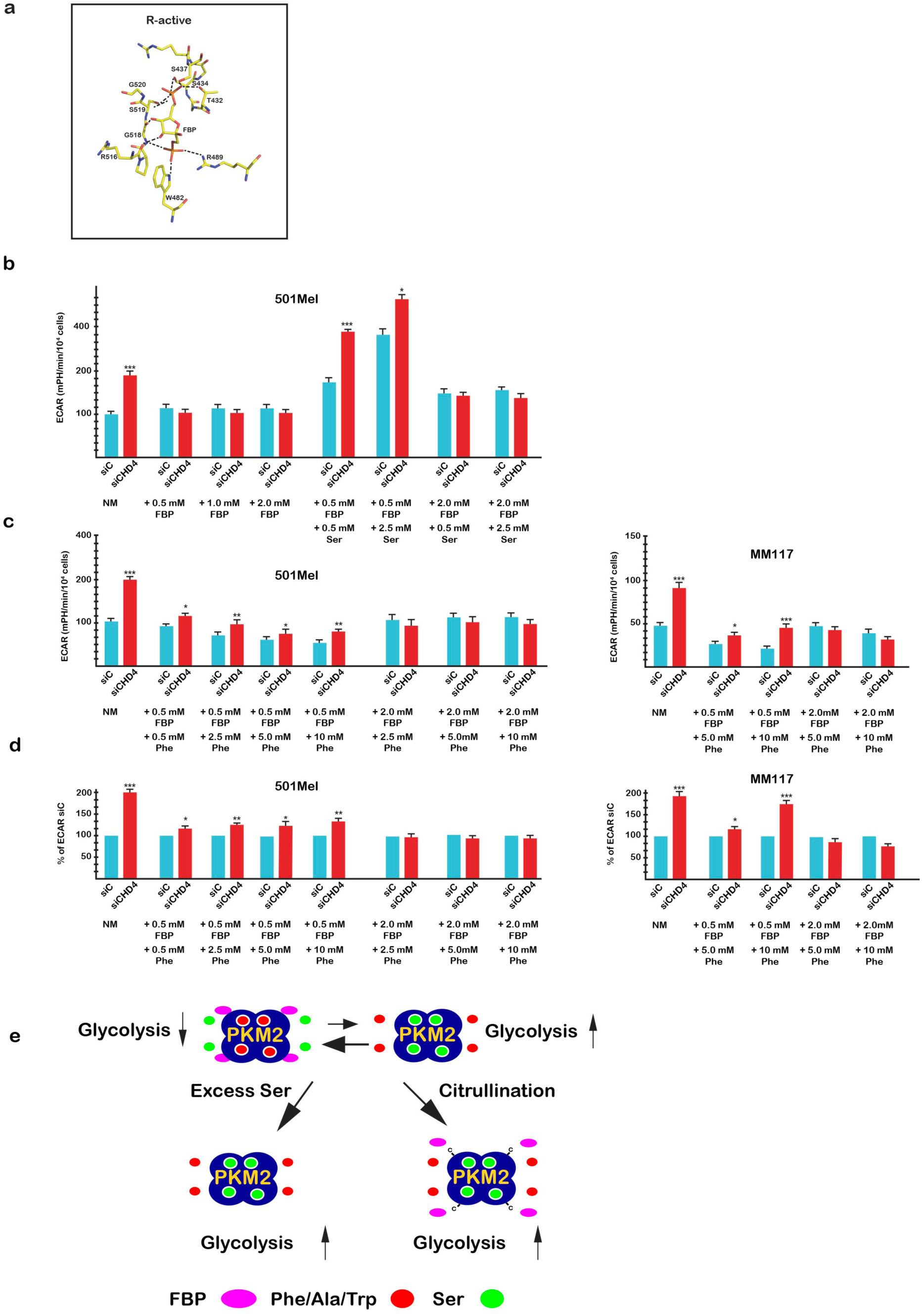
Effects of FBP and Ser on glycolysis. **a.** Close up view of FBP interactions with the R-active state illustrating the hydrogen bond between R489 and the 1’ phosphate of FBP as well as the network of hydrogen bonding with other residues. Salt bridges and hydrogen bonds are shown as dashed lines. For clarity, the side chain of K433 is not displayed. PDB data sets are as described in Supplementary Figure S3 **b**. ECAR values in presence of increasing exogenous FBP with or without exogenous Ser and siCHD4 silencing. NM = normal medium. **c.** ECAR values in presence of increasing exogenous Phe with or low or high exogenous FBP and siCHD4 silencing in 501Mel or MM117 cells as indicated. **d.** Effect of siCHD4 silencing on ECAR values in presence of increasing exogenous Phe with or low or high exogenous FBP expressed as a % of the siC control. N=6 with 6 technical replicates for each N. Unpaired t-test analysis were performed by Prism 5. P-values: *= p<0,05; **= p<0,01; ***= p<0,001. Data are mean ± SEM. **e.** A model for how citrullination affects PKM2 and glycolysis. Under basal conditions PKM2, represented as a tetramer, is in a dynamic equilibrium between a Ser bound form and a lower activity FBP bound form also in equilibrium with inhibitory amino acids. Increased Ser shifts the equilibrium to a Ser-bound form with higher activity due to mutually exclusive occupancy by Ser or Trp/Phe/Ala accounting for the observed increase in glycolysis. Citrullination diminishes FBP binding (R489<C represented by –C) alleviating its negative effect on Ser and shifts the mutually exclusive Ser vs Trp/Phe/Ala binding in favour of Ser. The net result is to promote a predominantly Ser-bound form accounting for the observed increased in glycolysis.

In agreement with this idea, increasing concentrations of exogenous FBP had little effect on basal glycolysis, but blocked stimulation by siCHD4 (Fig. 6b). Addition of exogenous Ser at low FBP concentration (0.5 mM) augmented basal glycolysis and re-established stimulation by siCHD4. In contrast, at higher FBP concentration (2.0 mM), no increase in basal or siCHD4-stimulated glycolysis was seen in presence of exogenous Ser. Increasing FBP therefore inhibited Ser and citrullination-dependent stimulation of glycolysis.

At low concentrations of exogenous FBP, increasing concentrations of exogenous Phe lowered basal glycolysis, whereas higher FBP concentrations overcame the Phe-induced repression (Fig. 6c-d), consistent with the known antagonistic effect of these ligands ^9, 10^. At low FBP concentrations and in presence of Phe, siCHD4 stimulated glycolysis, but to a lower level than seen in absence of FBP, whereas higher FBP concentrations blocked stimulation.

These observations indicated that increased FBP inhibited the ability of Ser to stimulate glycolysis under basal conditions and following citrullination. This is further supported by the observation that the ability of citrullination to overcome Phe inhibition through shifting the equilibrium towards Ser was also diminished by FBP. Together these observations support the idea that by disrupting its hydrogen bonding, R489 citrullination acted to lower FBP binding and its ability to inhibit Ser, while citrullination of R106 reduced inhibition by Phe/Ala/Trp shifting the equilibrium towards Ser. Through these two concerted events, citrullination therefore reprogramed PKM2 to be principally regulated by Ser (Fig. 6e).

## Discussion

### Citrullination; a novel regulator of PKM2 activity, glycolysis and cancer cell proliferation

Here we describe a regulatory pathway by which PKM2 citrullination regulates glycolysis and cancer cell proliferation. PKM2 is an allostatic regulator integrating a finely balanced feedback mechanism that modulates its activity over a wide range of absolute and relative amino acid concentrations ^9^.

FBP and Ser each stimulate PKM2 activity by stabilising the active R-state ^9^. Our data showed that exogenous FBP did not stimulate glycolysis in agreement with the report of Macpherson et al., ^10^ that intracellular FBP concentrations are sufficient to saturate PKM2. They also reported that FBP and Phe can simultaneously bind PKM2 with Phe preventing maximal activation of the FBP bound tetramer ^10^ maintaining PKM2 in a lower activity state as seen in tumours ^4^. Glycolysis was stimulated by exogenous Ser. Stabilisation of the active state by Ser, whose binding is mutually exclusive with Phe/Trp/Ala, would therefore lead to higher PKM2 activity compared to FBP. Ser depletion lowered basal glycolysis consistent with a dynamic equilibrium between a Ser-bound PKM2 and a less active FBP-Phe form that limits glycolysis and allows its dynamic regulation by changing this equilibrium (Fig. 6e). In agreement with this, exogenous Ser stimulated glycolysis, pushing the equilibrium towards Ser bound PKM2, whereas FBP that is already saturating and antagonized by Phe did not (Fig. 6e). Excess FBP did however counteract inhibition of glycolysis by exogenous Phe again consistent with their reported antagonism. Hence, while *in vitro* studies showed that FBP and Ser each stimulate PKM2 activity by stabilising the active R-state ^9^, our data show that FBP inhibited stimulation of glycolysis by Ser *in vivo*. This is unexpected given that FBP and Ser can bind PKM2 simultaneously at least under *in vitro* conditions used for crystallography. Our observations rather suggest that *in vivo*, the active R-state is stabilised by one or the other, but not by both simultaneously. How FBP antagonises stimulation by Ser remains to be determined.

The dynamic equilibrium regulating basal glycolysis can be upset by excess Ser but also by citrullination. R489 citrullination and diminished hydrogen bonding with FBP alleviated its negative effect on Ser-dependent stimulation of glycolysis. This effect is amplified by R106 citrullination that lowered PKM2 sensitivity to Trp/Ala/Phe shifting the equilibrium towards Ser. These two modifications could therefore act in concert to promote PKM2 regulation by Ser leading to increased glycolysis, an effect analogous to addition of excess Ser and in agreement with the observation that stimulation by citrullination required Ser (Fig. 6e). While we detected citrullination of multiple enzymes of the glycolytic pathway, our data on the effects of Ser, Phe/Ala/Trp and FBP on glycolysis all converge on PKM2 as being the central target. The role, if any, that citrullination exerts on the other glycolytic enzymes remains to be investigated.

Our data provide unique insight as to how conversion of arginine to citrulline impacts their key interactions. Unlike other post-translational modifications such as phosphorylation, or methylation, and to some extent acetylation, that often act positively to create new interactions with proteins the specifically recognize the modified amino acids, citrullination acts negatively due to loss of side chain charge and weakened hydrogen bonding ability. In the case of PKM2, our data illustrate how weakening of two interactions paradoxically translates into a positive reprograming and stimulation of glycolysis.

PKM2 has been shown to be regulated by other post-translational modifications, the best characterized of which are tyrosine phosphorylation ^12^, lysine acetylation on K305 ^32^ and K433 ^33^ and oxidation of C358 ^34, 35^. In each case, these modifications result in inhibition of PKM2 enzymatic activity. Moreover, most of the above studies concentrated on how post-translational modifications affected PKM2 activity after cell lysis or PKM2 immunoprecipitation overlooking that PKM2 activity in cells is regulated by a dynamic and complex crosstalk amongst its different ligands ^10^. Measuring glycolysis in the living cells was essential to assess how citrullination impacted cross-talk by multiple ligands to stimulate PKM2 and glycolysis. Citrullination is therefore a physiological mechanism that has an effect analogous to synthetic small molecules that increase PKM2 activity and stimulate excessive glycolysis resulting in Ser auxotrophy and reduced cell proliferation ^5, 6, 36, 37^.

Under most normal conditions, expression of PADI enzymes in general and PADI1 and PADI3 in particular is tightly regulated with low or no expression. There are however co-regulated in many human tumours. In addition, there are well documented situations where PADI enzyme expression is de-regulated. Expression of PADI enzymes can be induced under hypoxic conditions, for example in glioblastoma ^38^. In hypoxia, it has been shown that PKM2 undergoes proline hydroxylation and acts as a co-factor for HIF1A to increase expression of glycolytic enzymes ^11^. Our data further suggest that PADI1 and PADI3 expression in hypoxic tumour cells would stimulate glycolysis through PKM2 citrullination.

PADI enzyme expression is also de-regulated in pathological situations such as rheumatoid arthritis (RA) where the production of antibodies against aberrantly citrullinated proteins contributes to the chronic inflammatory state ^39–41^. Moreover, citrullination of glycolytic enzymes including PKM2 was observed in RA ^41^. The RA-associated environment is characterised by hypoxia and heterogeneous availability of nutrients, resembling that of some tumours ^42^. Thus, PADI1 and PADI3 expression and the subsequent PKM2 citrullination seen in RA may account for the increased glycolysis seen in activated RA-associated fibroblast-like synoviocytes, another hallmark of the disease ^42–44^.

In conclusion, we identify a novel pathway regulating melanoma cell proliferation where PADI1 and PADI3 citrullinate key arginines in PKM2 involved in its allosteric regulation to modulate glycolysis and cell proliferation. This pathway is shared in other cancer cells indicating a more general mechanism for regulating cell proliferation and may be active in other pathological contexts associated with increased glycolysis.

## Methods

A list of oligonucleotides, antibodies and resources can be found in Supplementary Dataset 3.

### Cell culture, siRNA silencing and expression vector transfection

Melanoma cell lines 501Mel and SK-Mel-28 were grown in RPMI 1640 medium supplemented with 10% foetal calf serum (FCS). MM074 and MM117 were grown in HAM-F10 medium supplemented with 10% FCS, 5.2 mM glutamax and 25 mM Hepes. Hermes-3A cell line was grown in RPMI 1640 medium (Sigma) supplemented with 10% FCS, 200nM TPA, 200pM cholera toxin, 10ng/ml human stem cell factor (Invitrogen) and 10 nM endothelin-1 (Bachem). HeLa cells were grown in Dulbecco’s modified Eagle’s medium supplemented with 10% FCS. SiHA cells were grown in EAGLE medium supplemented with 10% FCS, 0.1mM non-essential amino acids and 1mM sodium pyruvate. UOK cell lines were cultured in DMEM medium (4.5g/L glucose) supplemented with 10% heat-inactivated FCS and 0.1mM AANE.

SiRNA knockdown experiments were performed with the corresponding ON-TARGET-plus SMARTpools purchased from Dharmacon Inc. (Chicago, Il., USA). SiRNAs were transfected using Lipofectamine RNAiMax (Invitrogen, La Jolla, CA, USA) and cells were harvested 72 hours after. PADI1 and PADI3 expression vectors were transfected using X-tremeGENE^™^ 9 DNA Transfection Reagent (Sigma) for 48h. To assess clonogenic capacity, cells were counted and seeded in 6 well plates for 7 to 15 days.

### Proliferation, viability and senescence analyses by flow cytometry

To assess proliferation after siRNA treatment, cells were stained with Cell Trace Violet (Invitrogen) on the day of transfection. To assess cell viability, cells were harvested 72 hours after siRNA transfection and stained with Annexin-V (Biolegend) following manufacturer instructions. Cells were analysed on a LSRII Fortessa (BD Biosciences) and data were analysed using Flowjo software.

### ATP measurement

The concentration of ATP was determined 72h after siRNA transfection using the luminescent ATP detection system (Abcam, ab113849) following the manufacturer’s instructions.

### Protein extraction and Western blotting

Whole cell extracts were prepared by the standard freeze-thaw technique using LSDB 500 buffer (500 mM KCl, 25 mM Tris at pH 7.9, 10% glycerol (v/v), 0.05% NP-40 (v/v), 16mM DTT, and protease inhibitor cocktail). Cell lysates were subjected to SDS–polyacrylamide gel electrophoresis (SDS-PAGE) and proteins were transferred onto a nitrocellulose membrane. Membranes were incubated with primary antibodies in 5% dry fat milk and 0.01% Tween-20 overnight at 4 °C. The membrane was then incubated with HRP-conjugated secondary antibody (Jackson ImmunoResearch) for 1h at room temperature, and visualized using the ECL detection system (GE Healthcare). Antibodies: CHD3 abcam ab84528, CHD4 abcam ab72418, MITF abcam ab3201, SOX10, Abcam, ab155279.

### Generation of anti-Cit106 and anti-Cit246 PKM2 Antibodies

Antibodies against citrulline-containing peptides were raised in rabbits by the BioGenes company using the peptide sequences SFASDPILY-CIT-PVAVALDTKGGC and ASFI-Cit-KASDVHEVRKVLGEGGC for R106Cit and R246Cit respectively. Peptides were generated, quantified and confirmed by mass spectrometry by Genscript. Rabbits were immunized with carrier-conjugated peptide followed by three 3 booster injections after 14, 28 and 42 days. Affinity purification of antisera were performed by coupling the citrullinated peptides or their wild-type counterparts to SulfoLink Coupling Gel (PIERCE, 20401) agarose beads. The antisera were passed first through the column with citrullinated peptides and then through column with the wild-type peptide to remove residual antibodies recognizing the wild-type peptide.

### Immunoprecipitation and mass-spectrometry

Citrullinated proteins were immunoprecipitated from whole cell extracts with an anti-pan-citrulline antibody (Abcam, ab6464). Samples were concentrated on Amicon Ultra 0.5 mL columns (cutoff: 10 kDa, Millipore), resolved by SDS-PAGE and stained using the Silver 7 Quest kit (Invitrogen).

### Mass spectrometry and analysis

Mass-spectrometry was performed at the IGBMC proteomics platform (Strasbourg, France). Samples were reduced, alkylated and digested with LysC and trypsin at 37°C overnight. Peptides were then analyzed with an nanoLC-MS/MS system (Ultimate nano-LC and LTQ Velos ion trap, Thermo Scientific, San Jose Califronia). Briefly, peptides were separated on a C18 nano-column with a 1 to 30 % linear gradient of acetonitrile and analyzed in a TOP20 CID data-dependent MS method. Peptides were identified with SequestHT algorithm in Proteome Discoverer 2.2 (Thermo Fisher Scientific) *using* Human Swissprot database (20347 sequences). Precursor and fragment mass tolerance were set at 0.9 Da and 0.6 Da respectively. Trypsin was set as enzyme, and up to 2 missed cleavages were allowed. Oxidation (M) and Citrullination (R) were set as variable modifications, and Carbamidomethylation (C) as fixed modification.

Peptides were filtered with a 1 % FDR (false discovery rate) on peptides and proteins. For statistical analyses data was re-analysed using Perseus ^45^.

### Chromatin immunoprecipitation and sequencing

CHD4 ChIP experiments were performed on 0.4% Paraformaldehyde fixed and sonicated chromatin isolated from 501Mel cells according to standard protocols as previously described ^46^. MicroPlex Library Preparation kit v2 was used for ChIP-seq library preparation. The libraries were sequenced on Illumina Hiseq 4000 sequencer as Single-Read 50 base reads following Illumina’s instructions. Sequenced reads were mapped to the Homo sapiens genome assembly hg19 using Bowtie with the following arguments: -m 1 --strata --best -y -S -l 40 -p 2. After sequencing, peak detection was performed using the MACS software ^47^. Peaks were annotated with Homer (http://homer.salk.edu/homer/ngs/annotation.html) using the GTF from ENSEMBL v75. Peak intersections were computed using bedtools and Global Clustering was done using seqMINER. De novo motif discovery was performed using the MEME suite (meme-suite.org). Motif enrichment analyses were performed using in house algorithms as described in ^48^.

### RNA preparation, quantitative PCR and RNA-seq analysis

RNA isolation was performed according to standard procedure (Qiagen kit). qRT-PCR was carried out with SYBR Green I (Qiagen) and Multiscribe Reverse Transcriptase (Invitrogen) and monitored using a LightCycler 480 (Roche). RPLP0 gene expression was used to normalize the results. Primer sequences for each cDNA were designed using Primer3 Software and are available upon request. RNA-seq was performed essentially as previously described ^49^. Gene ontology analyses were performed with the Gene Set Enrichment Analysis software GSEA v3.0 using the hallmark gene sets of the Molecular Signatures Database v6.2 and the functional annotation clustering function of DAVID.

### Analysis of oxygen consumption rate (OCR) and glycolytic rate (ECAR) in living cells

The ECAR and OCR were measured in an XF96 extracellular analyzer (Seahorse Bioscience). A total of 20000 cells per well were seeded and transfected by siRNA or expression vector 72h and 24h hours respectively prior the experiment. The cells were incubated in a CO2-free incubator at 37°C and the medium was changed to XF base medium supplemented with 1mM pyruvate, 2 mM glutamine and 10mM glucose for an hour before measurement. For OCR profiling, cells were sequentially exposed to 2 µM oligomycin, 1 µM carbonyl cyanide-4-(trifluorome-thoxy) phenylhydrazone (FCCP), and 0.5 µM rotenone and antimycin A. For ECAR profiling, cells were sequentially exposed to 2 µM oligomycin and 150 mM 2-deoxyglucose (2-DG). After measurement, cells were washed with PBS, fixed with 3% PFA, permeabilized with 0.2% triton. Nuclei were counterstained with Dapi (1:500) and number of cells per well determined by the IGBMC High Throughput Cell-based Screening Facility (HTSF, Strasbourg). L-Phe (Sigma, P2126), L-Trp (Sigma, T0254), L-Ala (Sigma, A7627), L-Ser (Sigma, S4500) or D-FBP (Sigma, F6803) were added in the complete medium 24h for Serine and 6h for Trp/Phe/Ala/FBP and in the refreshed XF base medium prior the experiment.

## Acknowledgements

We thank, Dr Goncalo Castelo-Branco for the PADI3 expression vector, Drs JC. Marine and G. Ghanem for the MM117 and MM074 primary melanoma cells, Dr D. Bennet for the HERMES-3A line, all the staff of the IGBMC common facilities in particular the IGBMC mass spectrometry platform and the high throughput screening facility. This work was supported by institutional grants from the Centre National de la Recherche Scientifique, the Institut National de la Santé et de la Recherche Médicale, the Université de Strasbourg, the Association pour la Recherche contre le Cancer (CR, contract number PJA 20181208268), the Ligue Nationale contre le Cancer, the Institut National du Cancer, the ANR-10-LABX-0030-INRT French state fund through the Agence Nationale de la Recherche under the frame programme Investissements d’Avenir labelled ANR-10-IDEX-0002-02. The IGBMC high throughput sequencing facility is a member of the “France Génomique” consortium (ANR10-INBS-09-08). The mass spectrometry facility is supported by grants from the ARC foundation and from the Canceropole Grand Est. ID is an ‘équipe labellisée’ of the Ligue Nationale contre le Cancer. SC was supported by a fellowship from the Ligue Nationale contre le Cancer.

## Data availability

The CHD4 ChIP-seq and RNA-seq data described here have been deposited in GEO with the accession number GSE134850

## Author Contributions

SC performed ChIP-seq, RNA-seq, transfections and metabolism experiments, GD and GG performed bioinformatics analyses, LN performed and analysed mass-spectrometry experiments, SD constructed and provided PADI1 expression vector, CR performed structural analyses. SC, SD, CR and ID conceived the experiments, analysed the data and wrote the paper.

## Supplementary Figures and Legends.

**Supplementary Figure 1.**
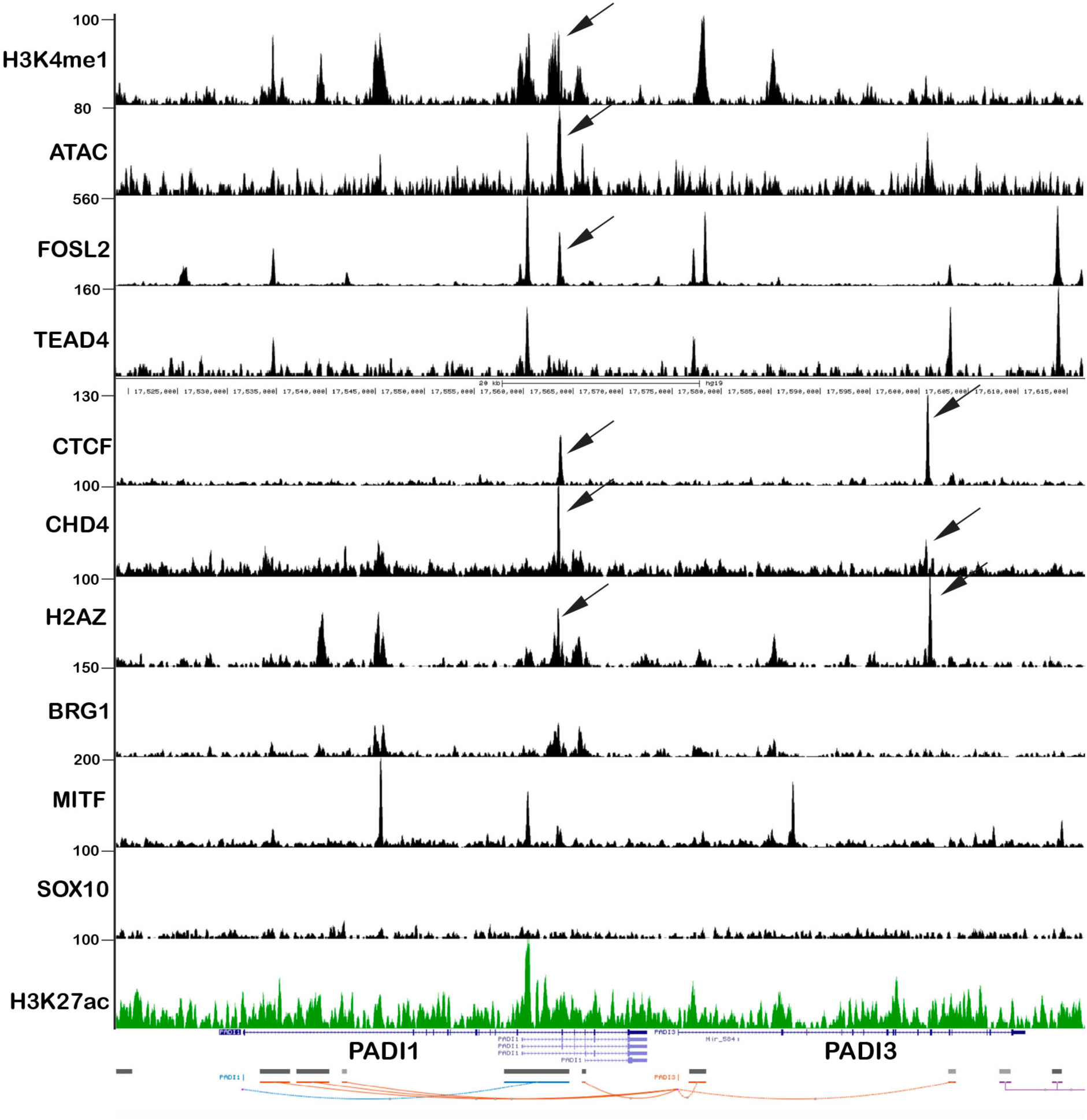
Regulation of the *PADI1*-*PADI3* locus. CHD4, CTCF and FOSL2 co-occupy a regulatory element at the *PADI1*-*PADI3* locus. Screenshot of UCSC genome browser at the *PADI1*-*PADI3* locus showing the indicated ChIP-seq data. Arrows highlight the putative cis-regulatory elements occupied by CTCF, FOSL1 and CHD4 and marked by ATAC-seq, H3K4me1, BRG1 and H2AZ. The following data sets were used: H3K4me1 GSM2476344; ATAC GSM2476338; FOSL2 GSM2842801; TEAD4 GSM2842802 ^50^; CHD4 this study. Other data are from Laurette et al., 2015 ^46^.

**Supplementary Figure 2.**
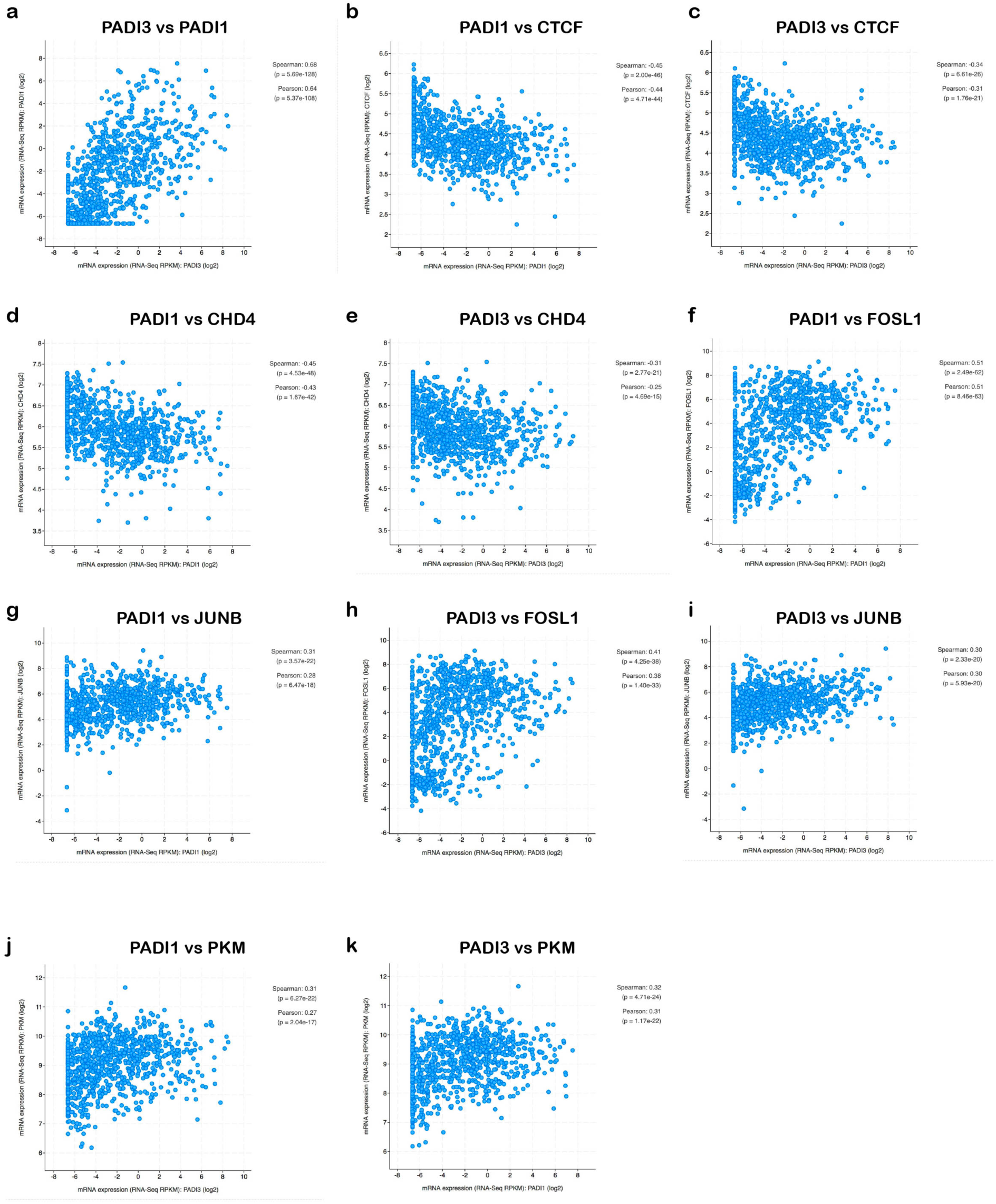
Analyses of gene expression in the Cancer Cell-Line Encyclopedia. Graphs shows that correlation of expression between the indicated genes with their Spearman and Pearson coefficients and p-values.

**Supplementary Figure 3.**
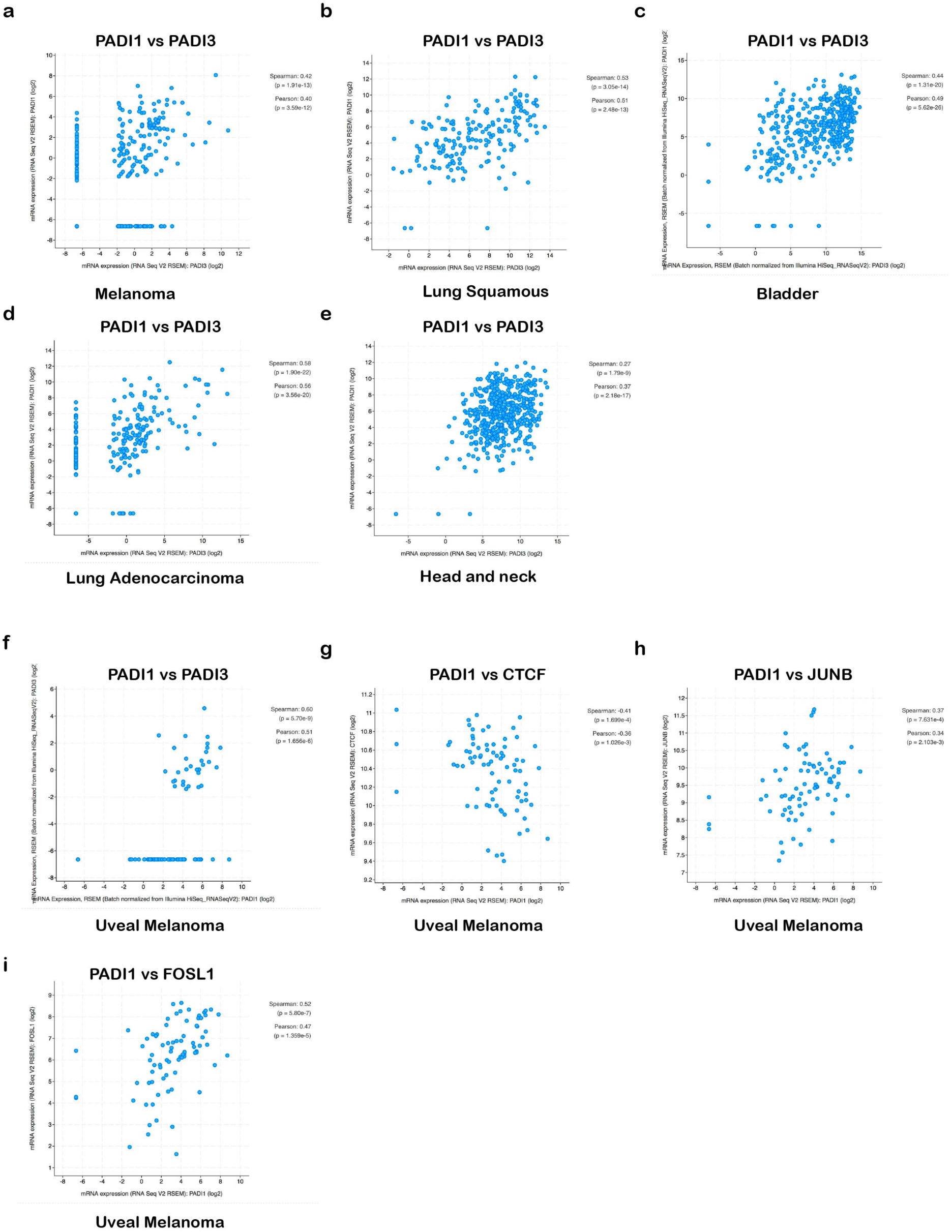
Analyses of *PADI1* and *PADI3* expression in The Cancer Cell Genome Atlas database. **a-f**. Graphs shows that correlation of *PADI1* and *PADI3* expression in the indicated tumour types with their Spearman and Pearson coefficients and p-values. **g-i** show correlation of *PADI1* with *CTCF*, *JUB* and *FOSL1* in uveal melanoma.

**Supplementary Figure 4.**
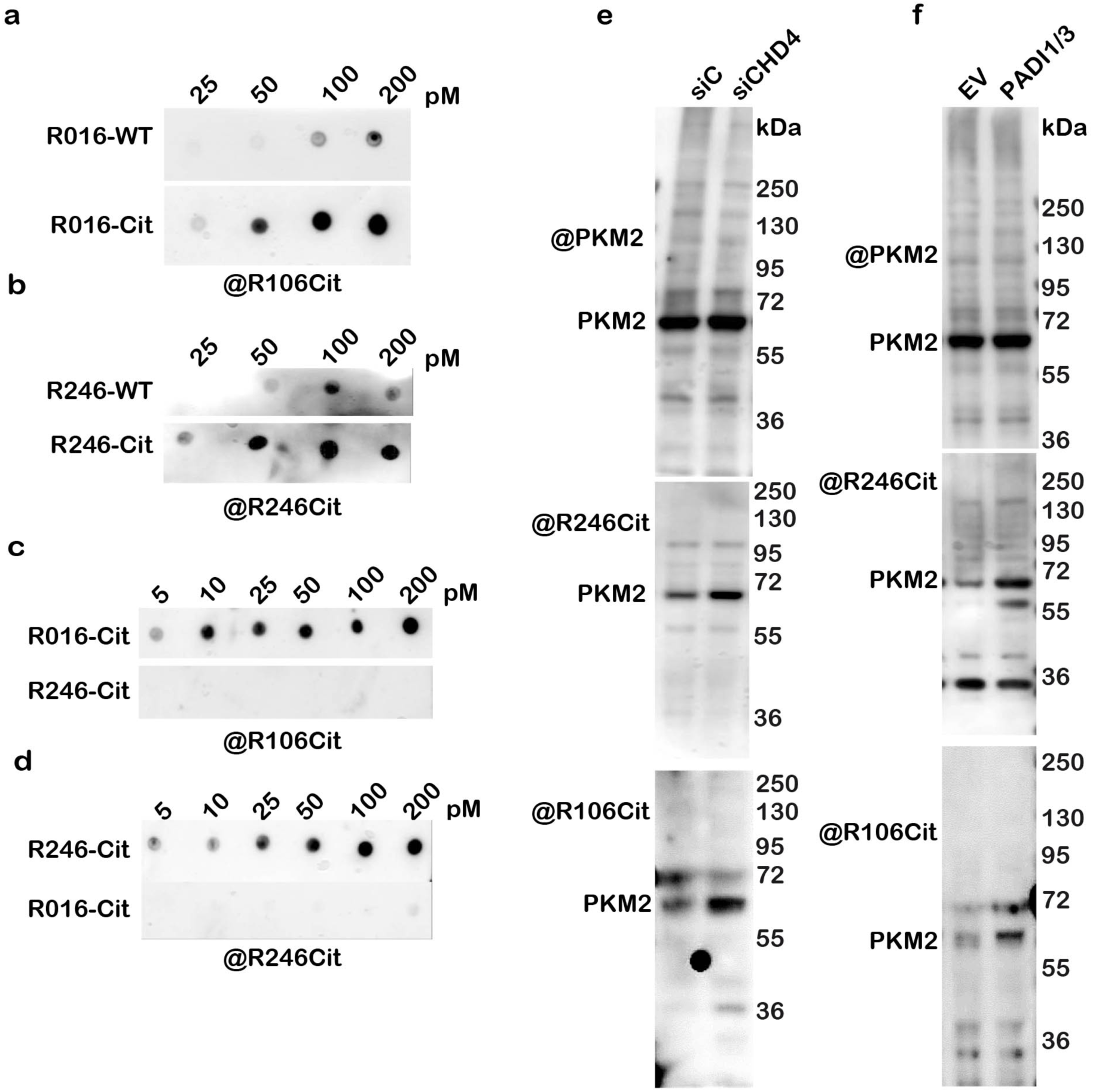
Enhanced citrullination of R106 and R246 following CHD4 silencing or ectopic PADI1 and PADI3 expression. **a-d**. Dot blots with the indicated amounts of wild-type peptides or equivalent peptides where R106 or R246 were replaced by citrulline. **e-f**. Immunoblots of cells transfected with the indicated siRNA or vectors. Cell extracts were loaded on three different gels and detected with commercial antibody against PKM2 to ensure equivalent quantities of PKM2 and with antibodies directed against the indicated citrullinated peptides. M shows the migration of the molecular mass standards.

**Supplementary Figure 5.**
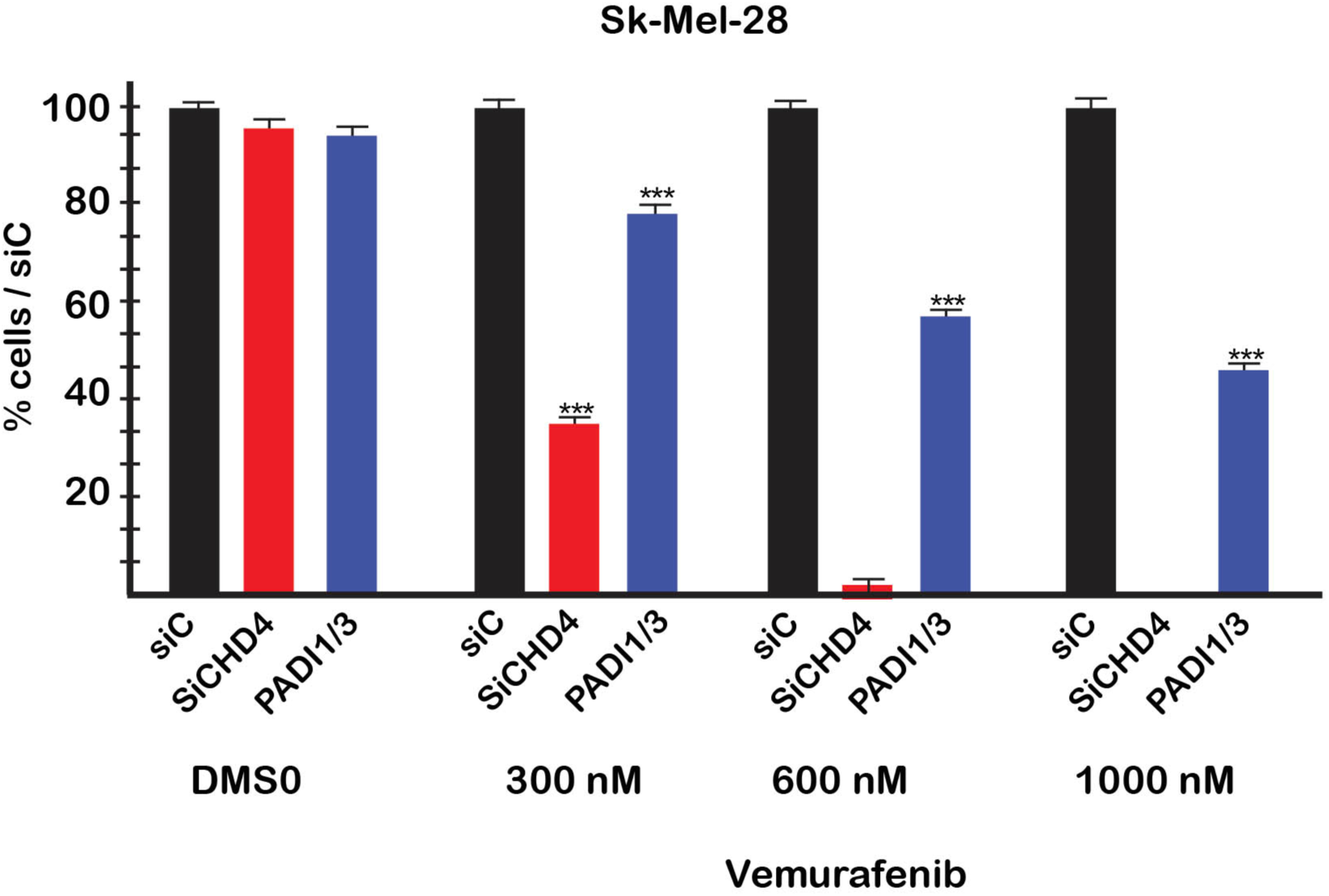
Citrullination modules sensitivity to BRAF inhibitor. Graph shows the % of surviving Sk-Mel28 cells compared to control 2 days after treatment with the indicated concentrations of vemurafenib.

**Supplementary Figure 6.**
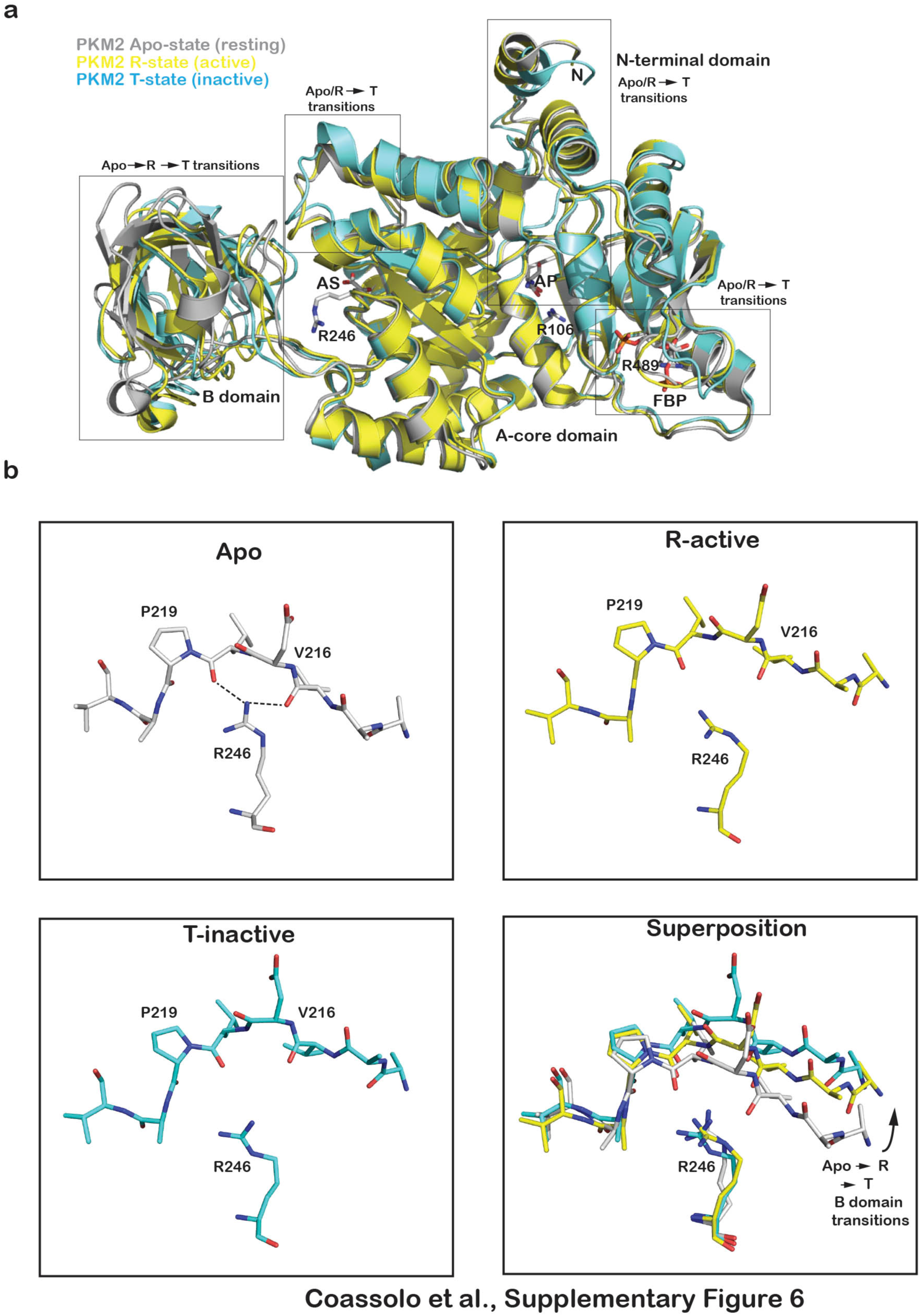
Locations and interactions of citrullinated arginines in PKM2. **a.** Ribbon representation of a PKM2 monomer in the apo resting state (grey; PDB 3SRH), the active R state (yellow; PDB 6GG6 with FBP and oxalate molecules from 3SRD) and the inactive T state (cyan; PDB 6GG4). The three citrullinated arginines (R106, R246 and R489), the free amino acids Serine and Phenylalanine, FBP and oxalate (surrogate of pyruvate to occupy the active site) are shown as sticks (carbon, grey; nitrogen, blue; oxygen, red; phosphorus, orange). AS, active site. AP, free amino acid binding pocket. The regions of PKM2 undergoing allosteric structural transitions between the three states are boxed. **b.** Closeup view of R246 interactions with the B domain in the Apo, R-active and T-inactive states along with a superposition of the three structures. Colour coding and representation of salt bridges/hydrogen bonds is as in panel a.

**Supplementary Dataset 1.** Summary of RNA-seq results following CHD3 or CHD4 silencing in 501Mel cells. Shown are gene names, description, fold change, p-value and adjusted p-value. As indicated, other pages on the spreadsheet show the ontology analyses of each gene set.

**Supplementary Dataset 2.** Proteins enriched after pan-citrulline immunoprecipitation from CHD4 silenced cells. Shown are accessions, gene names, gene descriptions, -Log P-values, differences (siCHD4-siCTRL), sum peptides scores, percentage of coverage, peptide number, PSM number, NSAF values (PSMs/protein length), unique peptide numbers, amino acid number and molecular mass.

**Supplementary Dataset 3.** A list of oligonucleotides, antibodies and resources.

